# PAR-4/LKB1 regulates intestinal cell number by restricting endoderm specification to the E lineage

**DOI:** 10.1101/2022.10.12.511872

**Authors:** Flora Demouchy, Ophélie Nicolle, Grégoire Michaux, Anne Pacquelet

## Abstract

The master kinase PAR-4/LKB1 appears as a major regulator of intestinal physiology. It is in particular mutated in the Peutz-Jeghers syndrome, an inherited disorder in which patients develop benign intestine polyps. Moreover, ectopic activation of PAR-4/LKB1 is sufficient to induce the polarized accumulation of apical and basolateral surface proteins and the formation of apical microvilli-like structures in intestinal epithelial cancer cell lines. In *C. elegans*, PAR-4 was shown to be required for the differentiation of intestinal cells. Here, we further examine the role of PAR-4 during intestinal development. We find that it is not required for the establishment of enterocyte polarity and plays only a minor role in brush border formation. By contrast, *par-4* mutants display severe deformations of the intestinal lumen as well as supernumerary intestinal cells, thereby revealing a novel function of PAR-4 in preventing intestinal hyperplasia. Importantly, we find that the ability of PAR-4 to control intestinal cell number does not involve the regulation of cell proliferation but is rather due to its ability to restrict the expression of intestinal cell fate factors to the E blastomere lineage. We therefore propose that PAR-4 is required to regulate *C. elegans* intestine specification.

## Introduction

The master kinase PAR-4/LKB1 is a notorious tumour suppressor, mutated in sporadic cancers as well as in the Peutz-Jeghers syndrome, an inherited disorder in which patients develop benign intestine hamartomatous polyps and a high frequency of malignant tumours (Alessi et al., 2006; Partanen et al., 2013). The exact origin of the intestinal polyps observed in the Peutz-Jeghers syndrome is not clear but may be linked to the function of LKB1 in intestinal stromal cells. Indeed, tissue-specific deletions of LKB1 in mice have shown that the loss of LKB1 in the sole intestinal epithelial cells does not induce polyp formation while LKB1 deletion in stromal cells, namely smooth muscle cells, fibroblasts and immune T cells, gives rise to polyposis (Katajisto et al., 2008; Ollila et al., 2018; Poffenberger et al., 2018). Abnormal production of interleukins in LKB1 deficient stromal cells may lead to increased JAK/STAT3 signalling and thereby to hyperproliferation of epithelial cells (Ollila et al., 2018; Poffenberger et al., 2018). Besides its role in preventing the formation of intestinal polyps, LKB1 has also been proposed to regulate intestinal polarity and brush border formation. In intestinal epithelial cancer cell lines, ectopic activation of PAR-4/LKB1 is sufficient to induce the polarized accumulation of typical apical and basolateral surface proteins and the formation of apical microvilli-like structures (Baas et al., 2004). It should however be noted that deletion of the LKB1 kinase domain in mouse intestinal cells does not seem to affect epithelial polarity (Shorning et al., 2009). Thus, although LKB1 appears as a major regulator of intestinal physiology, its exact *in vivo* functions remain to be elucidated. To further decipher the role of PAR-4/LKB1 during intestinal development, we took advantage of the *C. elegans* intestine. This very simple intestine constitutes an ideal model to characterize *in vivo* the different steps of intestinal development, including specification and differentiation, proliferation, polarization and formation of a brush border. The *C. elegans* intestine is composed of 20 cells, which form an antero-posterior tube surrounding the intestinal lumen. These 20 intestinal cells all derive from a single embryonic precursor cell, the E blastomere, which is born on the surface of the embryo at the 8-cell embryonic stage (Leung et al., 1999; Sulston et al., 1983). The two daughters of the E blastomere then migrate inside the embryo at the beginning of gastrulation, and undergo a series of stereotyped cell divisions, which give rise to the 20-cell stage intestine (Asan et al., 2016; Leung et al., 1999). The different intestinal development stages are named according to the number of E descendants present (E2, E4, E8, E16 and E20). Cell polarization occurs at the E16 stage (Achilleos et al., 2010; Leung et al., 1999; Totong et al., 2007). Shortly after, small gaps start to separate intestinal cells at the midline to eventually form the intestinal lumen (Asan et al., 2016; Bidaud-Meynard et al., 2021; Leung et al., 1999). Microvilli start to appear at the E20 stage, during the 1.5-fold embryonic developmental stage and form a regular brush border between the 2 and 3-fold stages (Asan et al., 2016; Bidaud-Meynard et al., 2021).

In 4-cell stage embryos, asymmetric cell division of the EMS cell gives rise to the E and MS blastomeres. The restricted expression of two GATA transcription factors, END-1 and END-3, in the E blastomere induces intestinal fate specification (Maduro et al., 2005a; Zhu et al., 1997, 1998) and expression of the differentiation GATA factors ELT-2 and ELT-7 (Fukushige et al., 1998; Maduro et al., 2005a; Sommermann et al., 2010; Wiesenfahrt et al., 2016; Zhu et al., 1998). *end-1* and *end-3* expression is induced by two other GATA factors, MED-1 and MED-2, which are themselves controlled by the bZIP-like factor SKN-1 (Maduro et al., 2005b, 2001). 50 to 80 % of embryos lacking SKN-1 or MED-1/2 do not form intestinal cells, reflecting the crucial role of the SKN-1 / MED-1/2 pathway for intestinal specification (Bowerman et al., 1992; Maduro et al., 2005b, 2001). However, the persistence of intestinal cells in a small percentage of these embryos also reveals the existence of parallel pathways, which can trigger intestinal specification. Consistently, further studies have shown that both the transcription factors POP-1/TCF and PAL-1 can also induce *end-1* and *end-3* expression in the E lineage (Maduro et al., 2005b). Moreover, while SKN-1 and MED-1/2 are expressed both in the E and MS blastomeres, their ability to induce *end-1* and *end-3* transcription is repressed in the MS cell. This repression is due to the strong nuclear accumulation of POP-1/TCF in the MS cell (Calvo et al., 2001; Lin et al., 1995; Maduro et al., 2005b; Shetty et al., 2005). In the E blastomere, Wnt signaling emanating from the P2 cell reduces the nuclear accumulation of POP-1/TCF, converts it to a transcription activator and allows both SKN-1 and POP-1 dependent *end-1* and *end-3* expression (Lin et al., 1995; Maduro et al., 2005b; Rocheleau et al., 1997; Shetty et al., 2005; Thorpe et al., 1997).

In *C. elegans*, PAR-4 plays a crucial role in the asymmetric localization of cell fate determinants in the early embryo (Kemphues et al., 1988; Tenlen et al., 2008) and is required for the differentiation of intestinal cells (Kemphues et al., 1988; Morton et al., 1992). In the one-cell embryo, it also moderately affects cortical polarity (Chartier et al., 2011; Hung and Kemphues, 1999) and controls cell cycle timing through the regulation of both CDC-25.1 and replication origins (Benkemoun et al., 2014; Rivers et al., 2008). In this work, we examine the role of PAR-4 during intestinal development. We show that PAR-4 is not required for epithelial polarity establishment and has a minor role in brush border formation. By contrast, severe deformations of the intestinal lumen are observed in *par-4* mutants. These deformations are associated with the presence of supernumerary intestinal cells. Importantly, we find that PAR-4 does not control on intestinal cell number by regulating cell proliferation but rather by restricting intestinal specification to the E blastomere.

## Results

### PAR-4 is dispensable for intestinal apico-basal polarity and microvilli formation in *C. elegans*

In *C. elegans*, a faint and uniform cortical PAR-4 immunostaining has been observed in one-cell embryos. As the embryos further divide, the cortical enrichment of PAR-4 becomes more robust, in particular at cell-cell boundaries (Watts et al., 2000). To further characterize PAR-4 expression pattern, we used CRISPR/Cas9 genome editing to endogenously label the two longest PAR-4 isoforms with the monomeric Neon Green (mNG) fluorophore. Consistent with a recent report (Roy et al., 2018), we observed that mNG::PAR-4 was ubiquitously expressed and cortically enriched in embryos, especially at cell-cell boundaries (Fig.1A). In enterocytes, mNG::PAR-4 localized both at the apical and basolateral cortex. During the early steps of intestine polarization, mNG::PAR-4 also localized on numerous subapical foci (Fig.1A). These foci were highly dynamic, suggesting that they may correspond to intracellular trafficking vesicles (Movie 1). Interestingly, we could distinguish two populations of foci: smaller foci moved in both directions all along lateral membranes while larger foci moved in the vicinity of the apical membrane.

**Figure 1:**
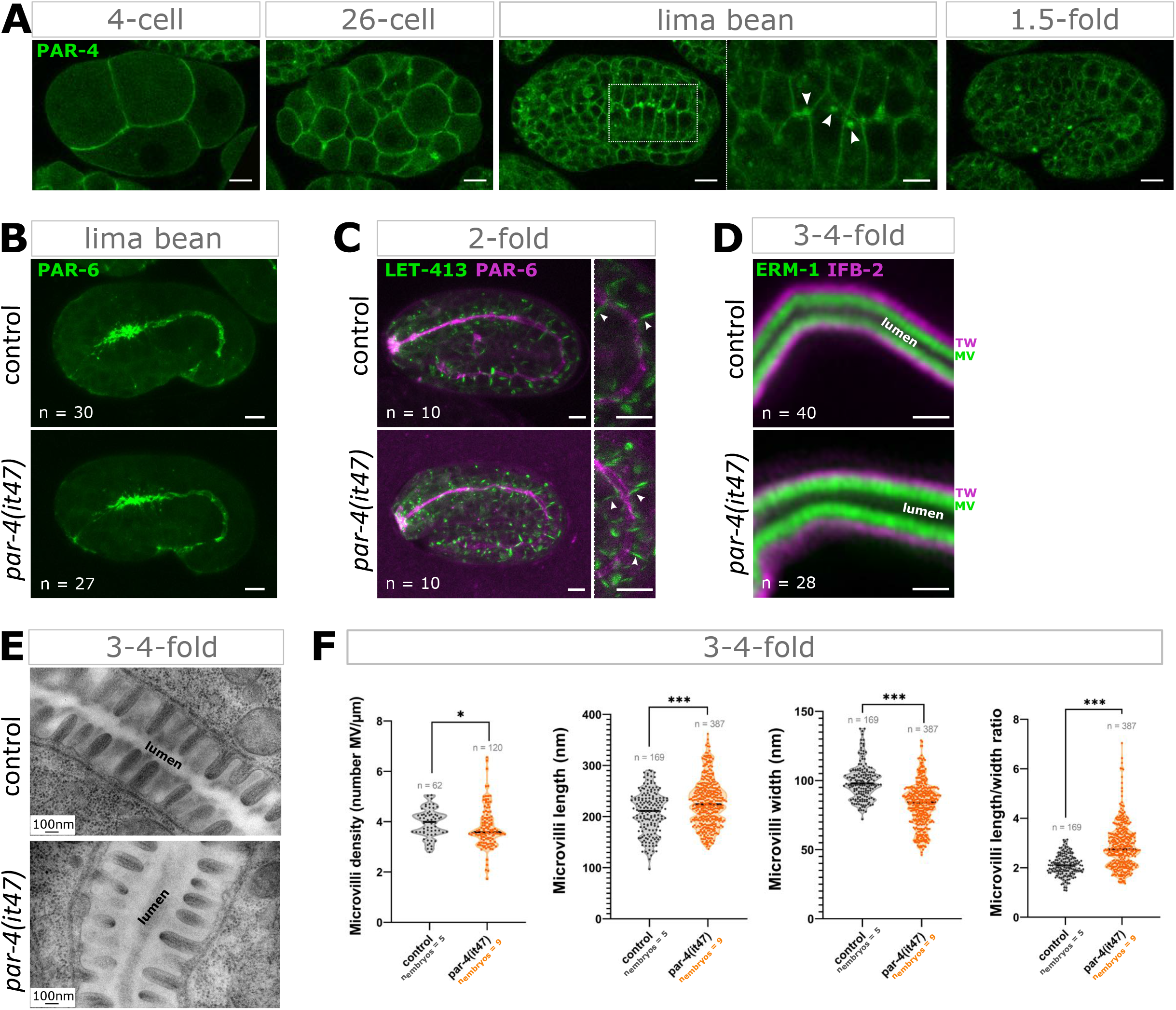
PAR-4 is not strictly required for intestinal cell polarity and microvilli formation in *C. elegans* embryos. **(A)** Confocal images of embryos of the indicated stages expressing endogenously tagged mNG::PAR-4. Scale bar: 6 μm. The higher magnification image at the lima bean stage shows the localization of mNG::PAR-4 in enterocytes, both on the apical and basolateral membranes and on subapical foci (arrowheads). Scale bar: 3 μm. **(B)** Maximum Z-projections from confocal images of lima bean control and *par-4(it47)* embryos expressing PAR-6::GFP. Scale bar: 5 μm. **(C)** Maximum Z-projections from confocal images of 2-fold control and *par-4(it47)* embryos co-expressing LET-413::mNG (green) and PAR-6::mKate2 (magenta). Zoomed-in images show basolateral LET-413::mNG signal (green, arrowheads) compared to apical PAR-6::mKate2 localization (magenta). Scale bar: 5 μm. **(D)** Super-resolution (SR) confocal microscopy images of 3-4-fold control and *par-4(it47)* embryos co-expressing ERM-1::mNG (green) and IFB-2::wScarlet (magenta). TW: terminal web. MV: microvilli. Scale bar: 1 μm. **(E)** TEM images of transversal sections showing the brush border in 3-4-fold control and *par-4(it47)* embryos. Scale bar: 100 nm. **(F)** Quantification of microvilli density, length, width and length/width ratio obtained from TEM images of 3-4-fold control and *par-4(it47)* embryos. Microvilli density corresponds to the number of microvilli per μm. Values were obtained from 5 control and 9 *par-4(it47)* embryos, measuring density in more than 7 areas and microvilli characteristics for more than 20 microvilli per embryo. Violin plots show all individual values (dots) and the mean (bold black line). They were obtained by pulling the data of all embryos. n indicates the total number of either areas or microvilli measured. ***P<0.001. Statistical significance was calculated using a Mann-Whitney test.

This localization prompted us to ask whether PAR-4 is necessary for *in vivo* polarization and brush border formation. Intestinal polarization starts with the apical accumulation of PAR-3/PAR-6/PKC-3 at the E16 stage, preceding embryonic elongation (Achilleos et al., 2010; Totong et al., 2007). Basolateral proteins such as LET-413^Scribble^ initially localize on all membranes before being excluded from the apical membrane at the beginning of embryonic elongation (Legouis et al., 2000; Pickett et al., 2021). Consistently, in elongated 2-fold embryos, we observed that endogenously tagged PAR-6::GFP and LET-413^Scribble^::mNG localized at the apical and basolateral membranes of enterocytes, respectively (Fig.1C). PAR-6::GFP already localized at the apical membrane prior to elongation, in early lima bean embryos (Fig.1B). To inactivate PAR-4 we used the *par-4(it47)* thermosensitive allele (Kemphues et al., 1988; Watts et al., 2000) and shifted the embryos at restrictive temperature (25°C) two hours before imaging. Inactivating PAR-4 two hours before the lima bean stage, i.e. prior to the beginning of enterocyte polarization, did not alter PAR-6 apical localization (n=27) (Fig.1B). Similarly, inactivating PAR-4 two hours before the 2-fold stage, i.e during the first step of apical polarization, before the apical exclusion of LET-413^Scribble^, did not prevent LET-413^Scribble^ from eventually being restricted to the basolateral membrane (n=10) (Fig.1C). Thus, PAR-4 does not seem to be required for the establishment of apico-basal polarity.

Once polarized, enterocytes assemble a regular array of microvilli which forms the brush border (Asan et al., 2016; Bidaud-Meynard et al., 2021; Leung et al., 1999). Observation of 3-4-fold embryos expressing endogenously tagged ezrin (ERM-1::mNG) and intermediate filament (IFB-2::wScarlet) by super-resolution confocal microscopy allowed us to distinguish regularly organized apical microvilli containing ezrin and the subapical terminal web (Fig.1D). Inactivating PAR-4 two hours before the 3-4-fold stage, i.e. during brush border formation, did not alter ERM-1 and IFB-2 localization and did not seem to prevent microvilli formation (n=28) (Fig.1D). To further characterize the effect of PAR-4 loss-of-function on brush border formation, we used transmission electron microscopy (TEM). At the 3 to 4-fold stage, control embryos displayed regular microvilli at the apical membrane of enterocytes (Fig.1E). Consistent with our super-resolution observations, we found that *par-4(it47ts)* embryos also developed an apical brush border (Fig.1E). Nonetheless, measurements of microvilli features, i.e their density, length and width, revealed mild defects of the brush border ultrastructure. Namely, microvilli density was slightly decreased while microvilli were slightly longer and thinner in *par-4(it47ts)* embryos compared to controls (Fig.1F). It should however be mentioned that our analysis also revealed rather strong inter-individual variations (Fig.S1A-C). Altogether, our observations demonstrate that PAR-4 is not required for intestinal cell polarity and has a rather minor role in microvilli formation in *C. elegans* embryos.

### PAR-4 regulates lumen architecture and intestinal cell number

While *par-4(it47)* embryos do not have strong polarity or brush border defects, we found that they displayed striking lumen deformations. In contrast with control embryos, which had a regular elliptical intestinal lumen, *par-4(it47)* mutant embryos exhibited severe deformations of the apical membrane (Fig.2A-B). At the 3-4-fold stage, such deformations were visible in 28% (n = 16/57) and 78% (n = 7/9) of *par-4(it47ts)* embryos observed either by confocal or transmission electron microscopy (TEM), respectively. Notably these defects were not observed in 2-fold embryos (n=20) (Fig.S2A). Importantly, we noticed that ERM-1 and IFB-2 correctly localized even in regions with strong lumen deformations (n=28) (Fig.S2B), thus confirming that apical membrane deformations are not linked to defects in apical polarity.

**Figure 2:**
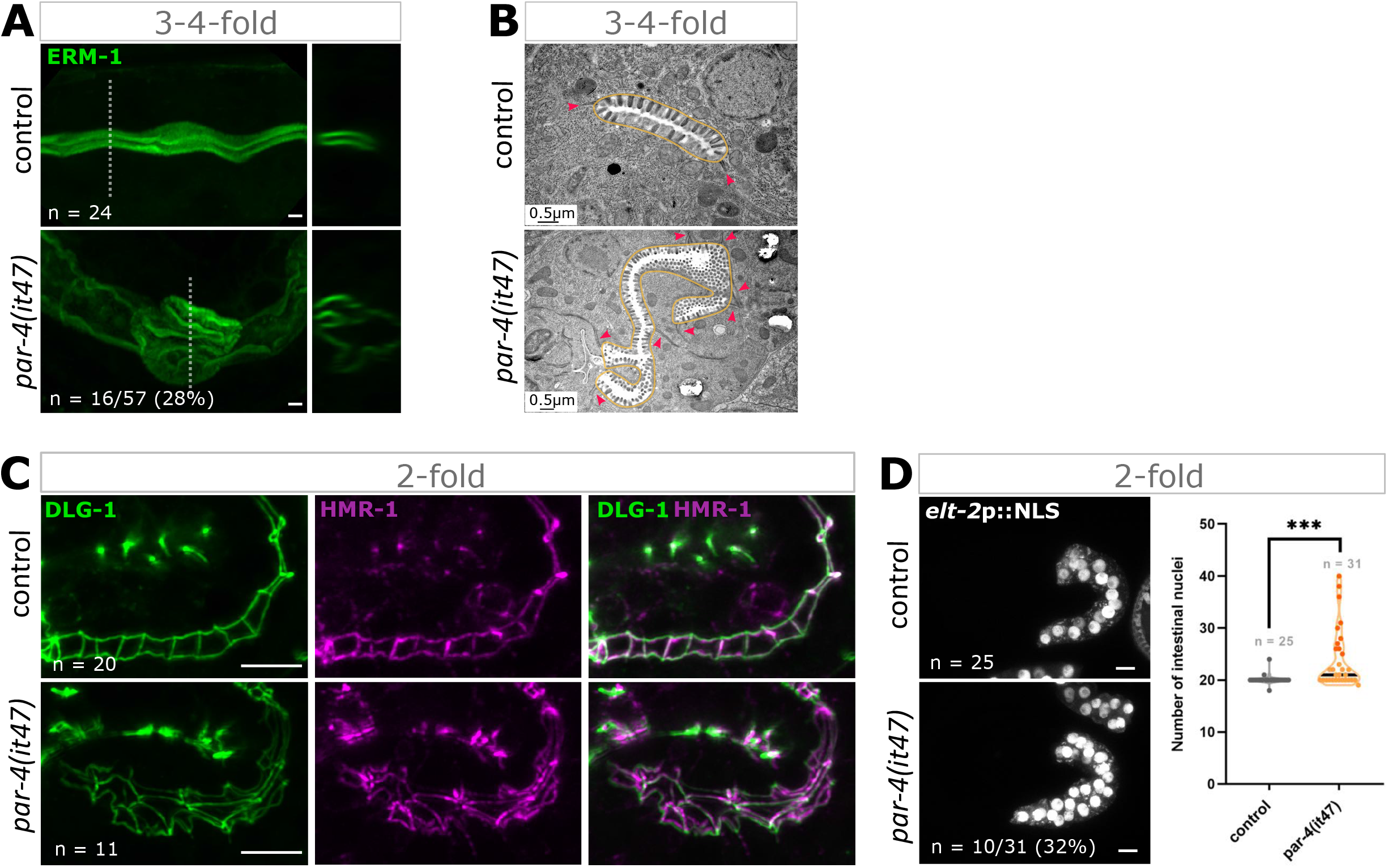
PAR-4 regulates lumen architecture and intestinal cell number. **(A)** 3-4-fold control and *par-4(it47)* embryos expressing ERM-1::mNG. Left and right panels show 3D reconstructions and orthogonal views, respectively. Dotted lines on the left panel indicate the plane used to show the orthogonal view on the right panel. Super-resolution images are shown for illustration, quantifications were obtained with embryos observed with a standard confocal microscope. Scale bar: 1 μm. **(B)** TEM images of transversal sections of 3-4-fold control and *par-4(it47)* embryos. The intestinal lumen is surrounded by a yellow line, apical junctions are indicated by red arrowheads. Scale bar: 0.5 μm. **(C)** Maximum Z-projections from SR confocal microscopy images of 2-fold control and *par-4(it47)* embryos co-expressing DLG-1::mNG (green) and HMR-1::mKate2 (magenta). Scale bar: 5 μm. **(D)** Maximum Z-projections from confocal images of 2-fold control and *par-4(it47)* embryos expressing *elt-2p*::NLS::GFP::LacZ and quantification of the number of intestinal nuclei in those embryos. Scale bar: 5 μm. Violin plots show all individual values (dots) and the mean (bold black line). Dark orange dots correspond to embryos with an excess of intestinal nuclei. ***P<0.001. Statistical significance was calculated using a Mann-Whitney test.

We next wondered whether these lumen deformations were associated with other defects in tissue organization and first examined apical junctions. *C. elegans* enterocytes have only one type of apical junction, the *C. elegans* apical junction (CeAJ), which is formed by two protein complexes, the cadherin/catenin complex (CCC) and the DLG-1/AJM-1 complex (DAC) (McMahon et al., 2001; Segbert et al., 2004). The CCC is composed of HMR-1, HMP-1 and HMP-2, the homologs of classical E-cadherin, α-catenin and β-catenin, respectively (Costa et al., 1998). The DAC localizes basally to the CCC and contains DLG-1, the homolog of Disc large (McMahon et al., 2001) and the non-conserved protein AJM-1 (Köppen et al., 2001). To test whether PAR-4 loss-of-function had an effect on intestinal CeAJs, we used strains expressing the endogenously tagged CeAJ components DLG-1::mNG, HMR-1::GFP or HMR-1::mKate2 and HMP-1::GFP (Heppert et al., 2018; Marston et al., 2016). DLG-1, HMR-1 and HMP-1 localized at CeAJs, both in control and *par-4(it47)* embryos (Fig.2C and S2C). However, additional CeAJs were visible in about one third of *par-4(it47)* embryos (Fig.2C, Fig.S2C-D). Contrary to lumen deformations, these supernumerary CeAJs were present both in 2-fold and 3-4-fold embryos (Fig.S2D). We next used TEM to complete our analysis. The *C. elegans* gut arranges into rings composed of two enterocytes, except the anterior ring, which is made of four cells. Thus, on TEM transversal sections, control embryos display two, occasionally four, electron dense structures corresponding to CeAJs, which maintain the cohesion between the two, or four, cells surrounding the lumen (Fig.2B). By contrast, more than two CeAJs were observed on several sections of all *par-4(it47)* embryos and sections with more than four CeAJs were found in 56% of *par-4(it47)* embryo (n = 5/9) (Fig.2B). Notably, we also found that junction length increased in all *par-4(it47)* embryos analyzed by TEM (n = 9) (Fig.S2E).

The presence of additional CeAJs prompted us to examine the number of intestinal cells present in *par-4(it47)* mutants. To this end, we used strains expressing an intestinal-specific nuclear marker, expressed under the control of the *elt-2* promoter, *elt-2p*::NLS::GFP::LacZ (Fukushige et al., 1998). While most control 2-fold embryos had 20 intestinal cells, 32% of *par-4(it47)* embryos (n = 10/31) had 25 or more enterocytes (Fig.2D), indicating that PAR-4 prevents the appearance of supernumerary intestinal cells. These observations were initially made after shifting *par-4(it47)* embryos two hours at restrictive temperature (25°C). We next wondered whether the timing of temperature shift could influence the number of intestinal cells. First, we counted intestinal cells in *par-4(it47)* embryos, which had been kept at permissive temperature (15°C) before observation and surprisingly found that 27% (n = 9/33) had an excess of enterocytes (Fig.S2F). This result could be explained by previous observations showing that the *par-4(it47)* allele is already partially inactivated at 15°C (Morton et al., 1992). However, the similarity between the phenotypes observed at 15°C or after two hours at 25°C also suggested that the presence of supernumerary cells was not due to the inactivation of PAR-4 in the two hours preceding the 2-fold stage but rather to an earlier effect of PAR-4. To test this hypothesis, we next tried to observe intestinal cells in *par-4(it47)* embryos after a longer shift at 25°C. Unfortunately, under those conditions, the *par-4(it47)* mutation induced pleiotropic embryonic defects, in particular an elongation arrest, precluding proper embryo staging and characterization.

### Lumen defects in *par-4* mutants are due do the presence of additional intestinal cells

Our observations revealed the existence of different gut phenotypes in *par-4(it47)* embryos: mild brush border defects, increased junction length, intestinal lumen deformations, additional apical junctions and excess of enterocytes. We wondered whether these phenotypes were due to a unique function or to several roles of PAR-4. We first asked whether the presence of supernumerary enterocytes could affect tissue organization and lead to the appearance of additional CeAJs and lumen deformations. To this end we observed embryos co-expressing an intestinal nuclear marker and the CeAJ marker HMR-1::mKate2 and found that 23% of *par-4(it47ts)* 2-fold embryos (n = 7/31) had extra intestinal nuclei. Importantly, all of them also displayed additional CeAJs (Fig.3A). Reciprocally, all embryos with additional CeAJs (n = 7/31) also had supernumerary intestinal cells. Moreover, observation of embryos co-expressing HMR-1::mKate2 and ERM-1::mNG showed that 61% of *par-4(it47)* 3-4-fold embryos (n = 25/41) displayed intestinal lumen deformations and 92% of those embryos (n = 23/25) also had additional CeAJs (Fig.3B). Reciprocally, 63% of *par-4(it47)* had additional CeAJs (n = 26/41) and 89% of them (n = 23/26) displayed lumen deformations. Similarly, the presence of additional CeAJs was associated with the presence of lumen deformations on TEM images (Fig.2B). We next used a plasma membrane marker specifically expressed in the intestine, *vha-6p*::PH::GFP, to be able to assess both lumen and enterocyte organization. In control 3-4-fold embryos, *vha-6p*::PH::GFP allowed us to distinguish the regular and elliptical shape of the lumen and the presence of two enterocytes in the ring surrounding the lumen (Fig.3C). By contrast, 18% of *par-4(it47)* embryos (n = 7/38) displayed intestinal lumen deformations (Fig.3C). Notably, all of them (n = 7/7) also had additional enterocytes surrounding the lumen (Fig.3C). Finally, our TEM analysis revealed the presence of more than 4 cells per intestinal ring on sections from 56 % of *par-4(it47)* embryos (n = 5/9). In all those cases, the lumen appeared deformed and additional CeAJs were observed (Fig.3D). Altogether, our observations reveal a strong correlation between the presence of extra intestinal cells, additional CeAJs and lumen deformation, suggesting that the presence of extra-intestinal cells could explain the presence of additional CeAJs and lumen deformations. By contrast, our TEM analysis did not reveal any obvious link between microvilli or CeAJs length defects and the number of additional CeAJs (Fig.S1A-C, S2E), suggesting that those latter defects may not be due to the presence of supernumerary enterocytes.

**Figure 3:**
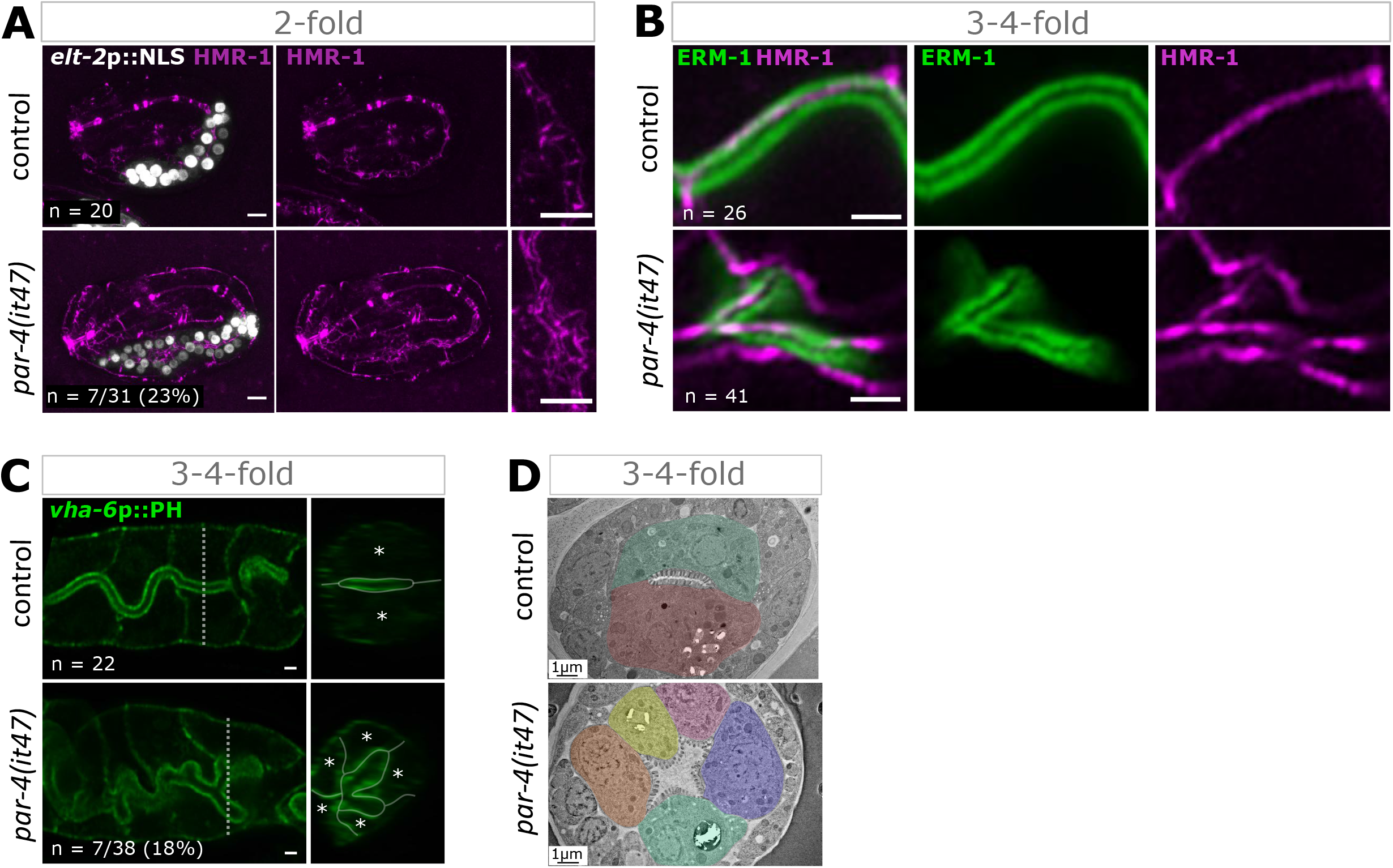
Defects in intestinal epithelium organization are due to the presence of additional cells. **(A)** Maximum Z-projections from confocal images of 2-fold control and *par-4(it47)* embryos co-expressing *elt-2p*::NLS::GFP::LacZ and HMR-1::mKate2. Zoomed-in images show high magnification of apical junctions (HMR-1::mKate2). Scale bar: 5 μm. **(B)** SR confocal microscopy images of 3-4-fold control and *par-4(it47)* embryos co-expressing ERM-1::mNG (green) and HMR-1::mKate2 (magenta). Scale bar: 1 μm. **(C)** Confocal images of 3-4-fold control and *par-4(it47)* embryos expressing *vha-6*p::PH::GFP. Left and right panels show longitudinal and orthogonal views, respectively. Dotted lines on the left panel indicate the plane used to show the orthogonal view on the right panel. White asterisks on orthogonal views indicate cells surrounding the lumen. Scale bar: 1 μm. **(D)** TEM images of transversal sections of 3-4-fold control and *par-4(it47)* embryos. Each intestinal cell is labelled by a different color. Scale bar: 1 μm.

### *cdc-25.1(gof)* mutants also display defects in lumen architecture

In order to confirm that the presence of extra intestinal cells is responsible for the presence of additional junctions and lumen deformations, we compared the different phenotypes of *par-4(it47)* embryos with those of *cdc-25.1* gain-of-function mutant embryos. Previous studies had indeed revealed the presence of supernumerary enterocytes in *cdc-25.1(rr31)* and *cdc-25.1(ij48)* gain-of-function mutants (Clucas et al., 2002; Kostić and Roy, 2002). Immunostaining of the junction component AJM-1 also suggested the presence of additional CeAJs in the intestine of *cdc-25.1(ij48)* embryos (Clucas et al., 2002). We first observed *cdc-25.1(rr31)* gain-of-function mutants co-expressing an intestinal nuclear marker and the CeAJ marker HMR-1::mKate2 and confirmed that all *cdc-25.1(rr31)* embryos had supernumerary enterocytes and additional CeAJs (n = 15) (Fig.S3A). We next characterized the lumen morphology of *cdc-25.1(rr31)* embryos expressing *vha-6p*::PH::GFP and observed that all of them (n = 18) displayed intestinal lumen deformations (Fig.4A). Similarly, TEM sections with more than four CeAJs and severe lumen deformations were observed in all analyzed *cdc-25.1(rr31)* embryos (n = 7) (Fig.4B). Consistent with previous observations (Choi et al., 2017), we found that additional enterocytes could be organized into rings composed of four (Fig.4A) or more cells (Fig.4B). Our TEM analysis also revealed that *cdc-25.1(rr31)* mutants displayed a slightly increased microvilli density and thicker microvilli while microvilli length and CeAJs length were not affected (Fig.S3B-D). Thus, while *par-4(it47)* and *cdc-25.1(rr31)* embryos share the same phenotypes in terms of lumen and junction organization, they do not show the same microvilli and junction length defects. Altogether, these observations are consistent with our hypothesis that additional junctions and lumen deformations are linked to the presence of extra intestinal cells but suggest that the defects of microvilli and CeAJs length observed in *par-4(it47)* mutant embryos are due to separate PAR-4 functions.

**Figure 4:**
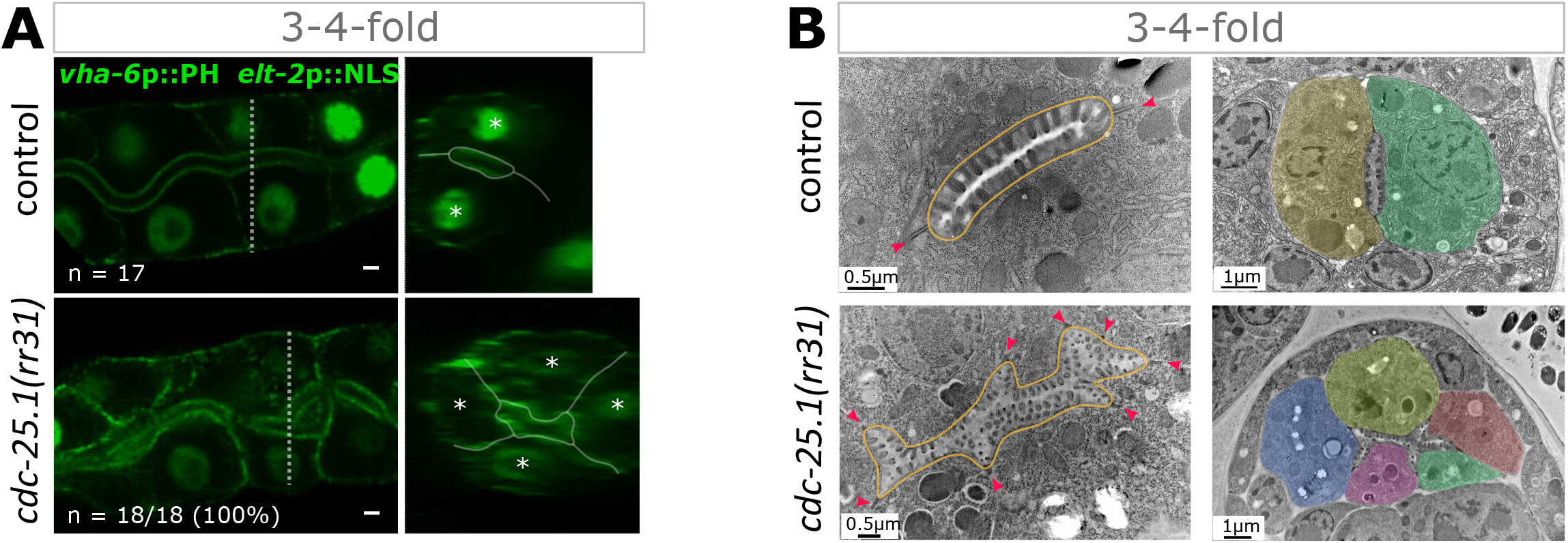
*cdc-25.1(gof)* mutants also display defects in lumen architecture. **(A)** Confocal images of 3-4-fold control and *cdc-25.1(rr31)* embryos expressing *vha-6*p::PH::GFP and *elt-2p*::NLS::GFP::LacZ. Left and right panels show longitudinal and orthogonal views, respectively. Dotted lines on the left panel indicate the plane used to show the orthogonal view on the right panel. White asterisks on orthogonal views indicate cells surrounding the lumen. Scale bar: 1 μm. **(B)** TEM images of 3-4-fold control and *cdc-25.1(rr31)* embryos. **(left)** The intestinal lumen is surrounded by a yellow line, apical junctions are indicated by red arrowheads. Scale bar: 0.5 μm. **(right)** Each intestinal cell is labelled by a different color. Scale bar: 1 μm.

### PAR-4 does not regulate cell proliferation in the E lineage

We next aimed at deciphering the mechanisms by which PAR-4 regulate intestinal cell number and first asked whether the supernumerary enterocytes observed in *par-4(it47)* embryos were due to additional divisions in the E lineage. In control embryos, the 20 enterocytes all emanate from the E blastomere, which undergoes 5 series of stereotyped divisions (Fig.5A) (Leung et al., 1999; Sulston et al., 1983). Acceleration of the cell cycle, as for instance reported in *cdc-25.1* gain-of-function mutant embryos, can lead to the appearance of an extra-round of divisions and supernumerary intestinal cells (Clucas et al., 2002; Kostić and Roy, 2002). We thus performed lineage experiments to determine whether PAR-4 also regulates the duration and number of intestinal cell divisions. To follow divisions, we used strains co-expressing two gut-specific nuclear markers expressed under the *end-1* or *elt-2* promoters, *end-1p*::GFP::H2B and *elt-2p*::NLS::GFP::LacZ (Fukushige et al., 1998; Shetty et al., 2005). The *end-1* promoter-driven endodermal reporter becomes visible in E2 cells before gastrulation whereas the *elt-2* promoter-driven intestinal marker appears during the E8 stage and persists afterwards (Fig.5B). In control embryos recorded at 25°C, it took around 3 hours between the appearance of the *end-1* promoter-driven signal in E2 cells and the E16 stage (Fig.5B, n = 13). Moreover, the transitions between E2 and E4, E4 and E8, and E8 and E16 stages took in average 36 (SD ± 3 min), 58 (SD ± 9 min) and 143 (SD ± 21 min) minutes, respectively (n = 13) (Fig.5C). As previously reported (Kostić and Roy, 2002), we found that intestinal cells divided faster in *cdc-25.1(rr31)* gain-of-function embryos: the transitions between E2 and E4, E4 and E8, and E8 and E16 stages took in average 28 (± SD 3 min), 29 (SD ± 3 min) and 100 (SD ± 37 min) minutes, respectively (n = 16) (Fig.5B-C). As a result, E descendants underwent an additional division and, after 3h at 25°C, the intestine was composed of 32 cells (Fig.5B). By contrast, no extra-division occurred in the E lineage of *par-4(it47)* embryos observed during 3h at 25°C (Fig.5B). Consistently, we did not observe any acceleration of the cell cycle as the transitions between E2 and E4, E4 and E8, and E8 and E16 took in average 38 (SD ± 6 min), 72 (SD ± 14 min) and 150 (SD ± 29 min) minutes, respectively (n = 19) (Fig.5C). Thus, unlike what happens in *cdc-25.1* gain-of-function mutants, the supernumerary intestinal cells observed *par-4(it47)* mutants do not result from an overall shortening of the E lineage cell cycles and from the appearance of an extra division between the E2 and E16 stage.

**Figure 5:**
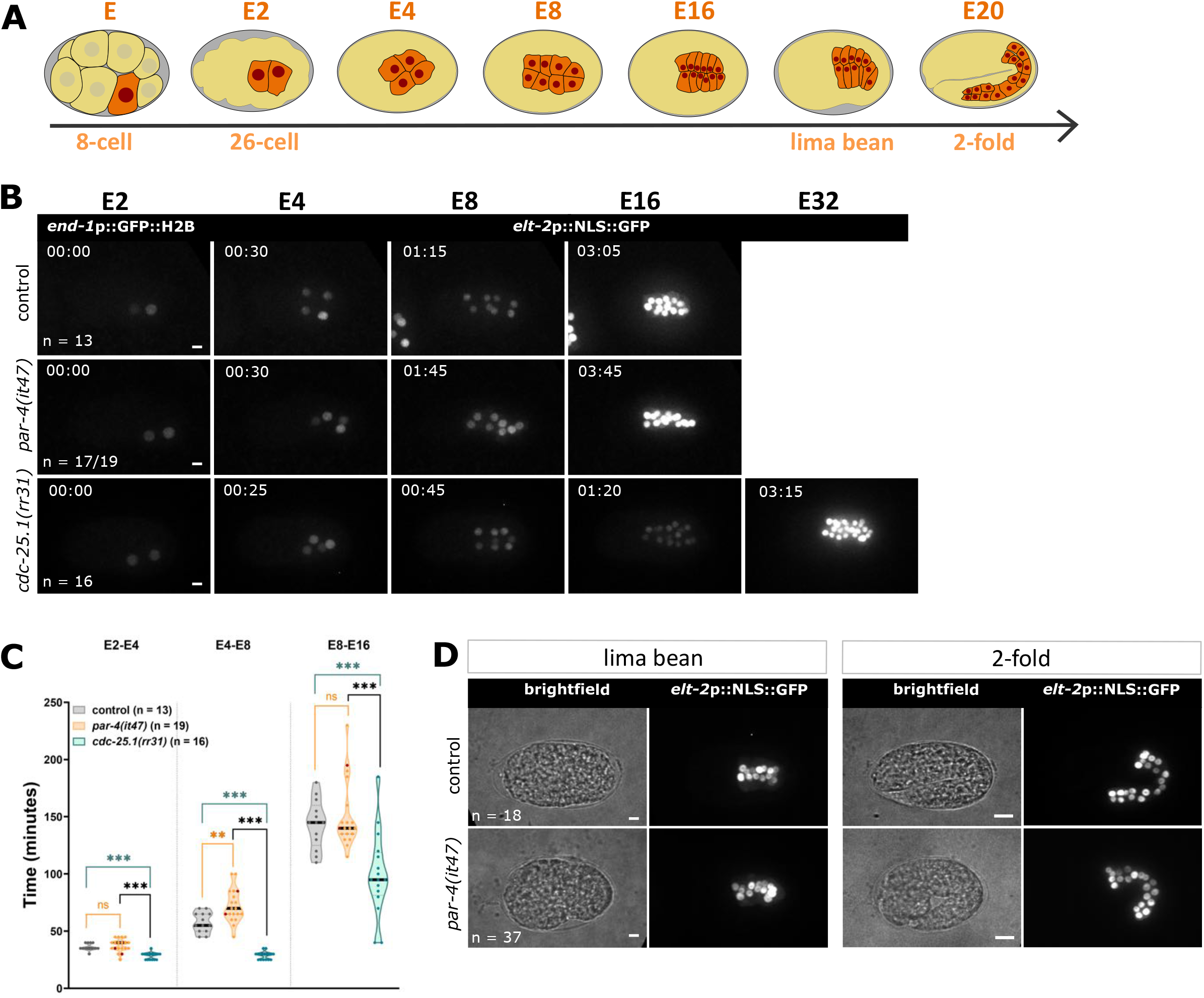
PAR-4 does not regulate cell proliferation in the E lineage. **(A)** Schematic representation of intestinal development stages during *C. elegans* embryogenesis. Enterocytes are generated by the E blastomere, which divides to give rise to E2, E4, E8 and E16 stages. The final E20 stage is reached at the 2-fold stage. Intestinal cells are represented in orange and their nuclei in dark red. **(B)** Maximum Z-projections from intestinal cell lineage movies in control, *par-4(it47)* and *cdc-25.1(rr31)* embryos. Embryos co-express two intestinal-specific nuclear markers: *end-1p*::GFP::H2B from E2 to E8 and *elt-2p*::NLS::GFP::LacZ from E8 to the end of intestinal cell divisions. In this experiment, 17/19 embryos showed such a phenotype. 2/19 embryos also exhibited ectopic expression of *end-1* outside the E lineage (not shown). Intestinal stages and corresponding times are indicated (in hour:min). Scale bar: 5μm. **(C)** Quantification of E2, E4 and E8 cell cycle lengths in control (n = 13), *par-4(it47)* (n = 19) and *cdc-25.1(rr31)* (n = 16) embryos. Violin plots show all individual values, plus the mean (bold black line). Dark red dots represent values in the two *par-4(it47)* embryos that ectopically express *end-1* signal in other cells than E descendants. **P<0.01, ***P<0.001, n.s: non-significant (P>0.05). Statistical significance was calculated using a Student’s *t*-test (E4-E8 control/*par-4(it47)* and E8-E16 control/*cdc-25.1(rr31)*) or a Mann-Whitney test (other comparisons). **(D)** Control and *par-4(it47)* embryos expressing *elt-2p*::NLS::GFP::LacZ were imaged at lima bean stage and one to two hours later. Embryo morphology is shown through the brightfield channel. Nuclei are shown with maximum Z-projections from confocal images. While all control embryos had started elongation, 7/37 *par-4(it47)* embryos did not elongate. Scale bar: 5μm.

Once embryos have reached the E16 stage, only four enterocytes of the E16 intestinal primordium, referred to as “star cells”, divide to obtain the final E20 stage (Sallee et al., 2021). These “star cells” usually divide between the late lima bean and the 1.8-fold stages ((Sallee et al., 2021) and our observations). During our long-lasting lineage experiments, we however failed to record this late division, including in control embryos. This prevented us from determining whether the extra intestinal cells observed in *par-4(it47)* mutants could result from the late and abnormal division of non “star-cells”. To circumvent this technical problem, we first imaged control and *par-4(i47)* embryos which were at the lima bean stage and had 16 intestinal nuclei. We next observed the same embryos between one and two hours later: at this time point, control embryos had mostly 20 intestinal cells (n = 15/18), occasionally 21 (n = 3/18) (Fig.5D). Similarly, *par-4(it47)* embryos mainly exhibited 20 intestinal nuclei (n = 32/37), occasionally 19 (n = 1/37), 21 (3/37) or 22 (1/37) (Fig.5D). They thus had normally completed their last intestinal cell division and did not undergo another round of division. Altogether, our results thus show that the presence of supernumerary intestinal cells in *par-4(it47)* mutants is not due to excessive cell proliferation in the E lineage. Surprisingly, we also observed that the intestine-specific degradation of PAR-4 did not affect intestinal cell number (Fig.S4A-B), suggesting that the presence of supernumerary intestinal cells in *par-4(it47)* mutants is not due the inactivation of PAR-4 in the E lineage but rather reveals a novel function of PAR-4 outside the E lineage.

### PAR-4 regulates intestinal cell number by restricting intestinal specification to the E lineage

While our lineage experiments showed that the descendants of the E blastomere divide normally in *par-4(it47)* embryos, we occasionally observed the expression of the *end-1* promoter-driven reporter in two cells outside the E lineage (n = 2/19). This ectopic expression of the *end-1* marker suggested that some cells that are not emanating from the E lineage adopt an intestinal-like fate and produce extra enterocytes. In these first lineage experiments, embryos were shifted at 25° just before imaging (see Table S2). We next tested whether an earlier shift would lead to the more frequent ectopic expression of the *end-1* marker and shifted the embryos at 25°C one hour before imaging. Under these conditions, we found that 53% of *par-4(it47)* embryos (n = 19/36) ectopically expressed *the end-1* reporter in one or two cells localized outside the E lineage (Fig.6A). This ectopic expression appeared at the E2 stage or when Ea and Ep divide (Fig.6A). Importantly, we found that the presence of these cells ectopically expressing *end-1* does not decrease cell cycle length in the E lineage (Fig.6B).

**Figure 6:**
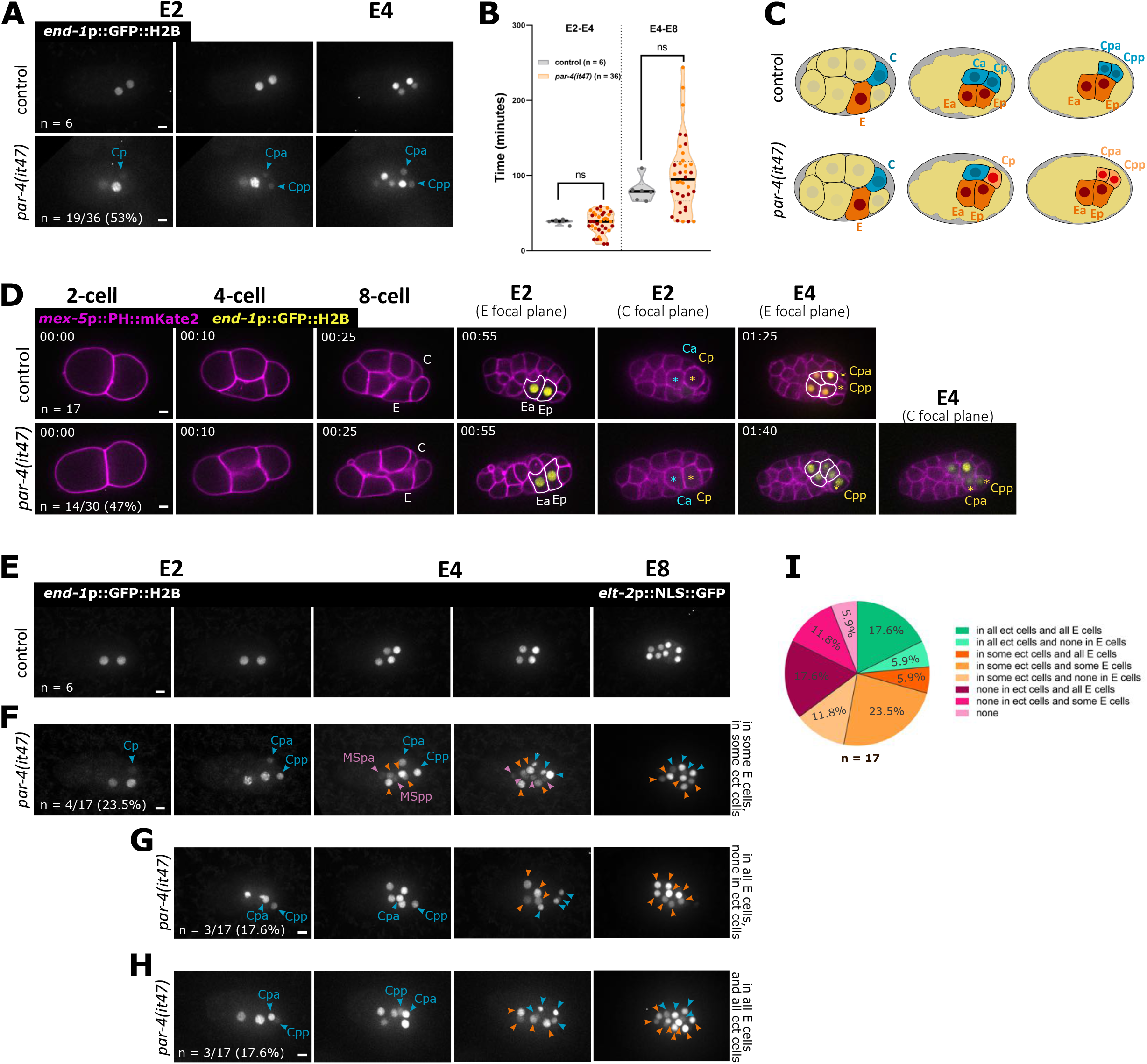
PAR-4 restricts intestinal specification to the E blastomere. **(A)** Maximum Z-projections from images of E2 and E4 stages from intestinal cell lineage movies in control and *par-4(it47)* embryos co-expressing *end-1*p::GFP::H2B and *elt-2p*::NLS::GFP::LacZ. 19/36 (53%) of *par-4(it47)* embryos present an ectopic *end-1* signal, most of them in the Cp lineage. Blue arrowheads indicate cells that ectopically express *end-1*p::GFP::H2B in the C lineage. **(B)** Quantification of E2 and E4 cell cycle lengths in control (n = 6) and *par-4(it47)* (n = 36) embryos. Violin plots show all individual values, plus the mean (bold black line). Dark red dots represent values in *par-4(it47)* embryos that ectopically express the *end-1* signal in other cells than E descendants. n.s: non-significant (P>0.05). Statistical significance was calculated using a Mann-Whitney test. **(C)** Schematic representation of 8-cell embryos and E2 stage during embryogenesis in control and *par-4(it47)* backgrounds. In control embryos, E and C descendants are drawn in orange and blue, respectively. In some *par-4(it47)* embryos, Cp adopts an intestinal-like fate by expressing *end-1*. **(D)** Images from movies in control and *par-4(it47)* embryos that co-express *end-1p*::GFP::H2B (yellow) and a *mex-5p*::PH::mKate2 membrane marker (magenta). Intestinal stages and corresponding times are indicated (hour:min). Ea and Ep and their daughters are marked with white lines. Ca is marked with a blue asterisk and Cp and its daughters with a yellow asterisk. Scale bar: 5μm. **(E-H)** Maximum Z-projections from images from intestinal cell lineage movies from E2 to E8 stages in control **(E)** and *par-4(it47)* embryos **(F-H)**. Embryos co-express *end-1p*::GFP::H2B and *elt-2p*::NLS::GFP::LacZ. The *elt-2p*::NLS::GFP::LacZ signal starts to be visible at the E8 stage. On those images, signal intensities were adjusted to optimize the visualization of intestinal nuclei at all stages. On non adjusted images, the strong intensity of the *elt-2p*::NLS::GFP::LacZ signal allows it to be easily distinguished from the much weaker *end-1p*::GFP::H2B signal. Orange arrowheads indicate E descendants. Blue and pink arrowheads indicate cells that ectopically express *end-1*p::GFP::H2B in the Cp and the MSp lineage, respectively. In *par-4(it47)* embryos exhibiting ectopic *end-1* signal, *elt-2* can for example be expressed in some E cells and in some ectopic cells (F), in all E descendants but not in ectopic cells (G) or in all E cells and ectopic cells (H). In (F), 5 descendants of Cp are present on the last image, only 4 express *elt-2* and are indicated with blue arrowheads, 8 descendants of E are present, only 5 express *elt-2* and are indicated with orange arrowheads. In (H), 4 descendants of Cp and 8 descendants of E are present, all express *elt-2*. Scale bar: 5μm. **(I)** Pie chart representing the distribution of *elt-2* expression in *par-4(it47)* embryos that exhibit ectopic *end-1* signal. Each color corresponds to a distinct phenotype. Percentages of each phenotype are indicated.

According to their position within the embryo, most of E-like cells expressing *end-1* in *par-4(it47)* embryos seem to arise from the C lineage (Fig.6A). With the help of the brightfield channel we could follow cell lineages and unambiguously identify the origin of these E-like cells in 17 embryos. We found that the *end-1* reporter was mainly expressed in the posterior daughter of the C blastomere, called Cp, or its descendants, Cpa and Cpp (n = 15/17) (Fig.6A,C, Fig.S5) and occasionally both in Ca and Cp (2/17). In 3/17 embryos, the *end-1* reporter was expressed both in the C lineage and in the MSp lineage (Fig.6F). To confirm these results, we followed embryonic cell divisions by imaging embryos co-expressing the *end-1* reporter and a ubiquitously expressed membrane marker (Fig.6D). In this experiment, 47% of *par-4(it47)* embryos (n = 14/30) showed ectopic expression of the *end-1* marker. This mostly occurred within the C lineage (n = 10/14), especially in Cp or its descendants Cpa and Cpp (n = 9/10) and occasionally both in Ca and Cp (n = 1/10). In a few embryos (n = 4/14), the *end-1* marker was expressed only in MSp descendants. Our data thus suggests that PAR-4 restricts intestinal specification to the E lineage by preventing *end-1* expression in the C and MS blastomeres.

We then wondered whether the ectopic expression of *end-1* in cells localized outside the E lineage is sufficient to induce intestinal differentiation in those cells. To address this question, we analyzed the expression of the *elt-2-*promoter driven reporter in *par-4(it47)* embryos displaying ectopic *end-1* expression. All control embryos (n = 6), as well as all *par-4(it47)* embryos without *end-1* ectopic expression (n = 17), expressed the *elt-2* reporter in all E descendants (Fig.6E). Among the *par-4(it47)* embryos ectopically expressing *end-1* in the C lineage, 65% (n=11/17) also expressed the *elt-2* reporter in at least some descendants of the C lineage. The number of C descendants expressing the *elt-2* reporter was variable, ranging from no cell to all cells ectopically expressing *end-1* (Fig.6F-I). Notably, in those embryos, there was also substantial variability in the number of E descendants expressing *elt-2* (Fig.6F-I). Unlike the C lineage, the descendants of MSp ectopically expressing the *end-1* marker never turned on the *elt-2* signal (Fig.6F). This high variability in the expression of the *elt-2* reporter may be due to a more variable *end-1* expression in *par-4(it47)* embryos, both inside and outside the E lineage.

Altogether, our observations demonstrate that PAR-4 regulates the number of enterocytes without controlling cell proliferation in the E lineage but rather but by preventing the C blastomere from adopting an intestinal fate. When PAR-4 is inactivated, *end-1* is ectopically expressed in the C lineage where it is sufficient to induce the ectopic expression of *elt*-2, thereby leading to the differentiation of supernumerary intestinal cells.

## Discussion

In this paper, we have shown that PAR-4 is not required for the establishment of apico-basal polarity during *C. elegans* intestinal development and has a minor role during brush border formation. By contrast, PAR-4 is essential to prevent intestinal hyperplasia. PAR-4 inactivation indeed results in the presence of supernumerary intestinal cells and severe lumen deformations. This novel function of PAR-4 appears to be independent from cell proliferation regulation and involves restriction of intestinal specification to the E lineage.

Our endogenously expressed mNG::PAR-4 construct confirms that PAR-4 is ubiquitously expressed in *C. elegans* embryos (Roy et al., 2018; Watts et al., 2000). It is enriched at the cell cortex, especially at cell-cell boundaries. As previously described (Watts et al., 2000), it is not polarized along the antero-posterior axis in early embryos. It also localizes to both the apical and lateral membranes in the intestinal epithelium. Besides its cortical localization, PAR-4 also localizes to dynamic foci in polarizing enterocytes. Our observations suggest the existence of two types of PAR-4 foci, small ones, which move along the lateral membranes and larger ones close to the apical membrane. Whether these two populations of PAR-4-positive foci have distinct functions and are associated with different types of trafficking vesicles remains to be determined.

Despite its intriguing dynamic localization in polarizing enterocytes, PAR-4 appears not to be required for the establishment of intestinal apico-basal polarity in *C. elegans*. We can not exclude that PAR-4 plays a minor, redundant role during polarization but it is clearly not an essential player in that process. Our results contrast with studies showing that ectopic activation of LKB1 is sufficient to establish polarity in isolated intestinal cancer cells (Baas et al., 2004). They are however consistent with the absence of obvious polarity defects in mice intestinal cells lacking LKB1 activity (Shorning et al., 2009). PAR-4/LKB1 has been shown to modulate the asymmetric localization of PAR-3, PAR-6 and aPKC in several other *in vivo* systems, including in epithelial cells (e.g. (Amin et al., 2009; Bonaccorsi et al., 2007; Chartier et al., 2011; Lee et al., 2007; Martin and St Johnston, 2003)). Many cell types can nevertheless apparently polarize normally in its absence (e.g. (Amin et al., 2009; Krawchuk et al., 2015), underlining that PAR-4/LKB1 is far from systematically having a crucial role in polarity establishment. Its requirement appears to be extremely sensitive to the cell context and may depend on the robustness of other polarization mechanisms at play in each cell type.

Our results show that PAR-4 is not strictly required for microvilli formation in *C. elegans* embryos but contributes to the formation of very typical microvilli. In intestinal cell lines, ectopic activation of LKB1 triggers the formation of microvilli-like structures through the activation of a signalling cascade involving the small GTPase Rap2 and ezrin phosphorylation (Baas et al., 2004; Gloerich et al., 2012; ten Klooster et al., 2009). In *C. elegans*, ERM-1 phosphorylation prevents the basolateral accumulation of ERM-1 in embryonic intestinal cells (Ramalho et al., 2020). The normal apical localization of ERM-1 that we observed in *par-4* embryos thus strongly suggests that ERM-1 phosphorylation is not affected in this context. Moreover, our TEM observations did not reveal microvilli defects in *rap-2(gk11)* mutant larvae (O. Nicolle, unpublished observations). It is thus unlikely that PAR-4 regulates microvilli growth through the RAP-2/ERM-1 pathway in *C. elegans*.

Besides its effect on microvilli morphology, we find that PAR-4 regulates CeAJ length. Interestingly, drosophila *lkb1(-)* photoreceptor cells also exhibit longer adherens junctions (Amin et al., 2009). Those are also frequently fragmented, a phenotype that we did not observe in the intestine of *par-4 C. elegans* mutants. In *C. elegans*, LET-413^Scribble^ has been proposed to trigger CeAJs compaction and *let-413* mutants display extended and fragmented intestinal CeAJs (Legouis et al., 2000; McMahon et al., 2001). The longer CeAJs observed in *par-4(it47)* embryos could therefore in principle result from reduced LET-413 activity. However, loss of LET-413 also results in the strong mislocalization of the apical proteins PAR-3 and PAR-6 (McMahon et al., 2001). The absence of polarity defects in *par-4(it47)* embryos argues against a role of PAR-4 in regulating LET-413 activity.

Intestinal hyperplasia is the most striking phenotype that we observe in *par-4* embryos. In 2-cell *C. elegans* embryos, *par-4* mutants show an acceleration of the cell cycle in the P1 blastomere (Morton et al., 1992). This effect can be explained by the ability of PAR-4 and its effector PAR-1 to inhibit CDC-25.1 nuclear accumulation as well as DNA replication in P1 (Benkemoun et al., 2014; Rivers et al., 2008). Moreover, stabilization of CDC-25.1 in the intestine leads to cell cycle acceleration and extra enterocytes divisions (Clucas et al., 2002; Kostić and Roy, 2002). However, in the intestine, *par-4* mutants exhibit neither cell cycle acceleration nor extra divisions, suggesting that PAR-4 does not regulate CDC-25.1 in that context and that intestinal hyperplasia is not linked to excessive cell proliferation.

Rather we find that supernumerary cells arise from specification defects and are due to the abnormal expression of the intestinal specification gene *end-1* in the C lineage. In control embryos, expression of *end-1* in the E blastomere is mostly activated by SKN-1 but also by POP-1/TCF and PAL-1 (Maduro et al., 2005b). In the C lineage, SKN-1 accumulation is limited by the kinase GSK-3: depletion of GSK-3 leads to SKN-1 stabilization and transformation of C descendants into E-like cells (Maduro et al., 2001; Schlesinger et al., 1999; Shirayama et al., 2006). While PAL-1 is present and activates *end-1* expression in E cells, it is expressed at higher levels in C descendants where it promotes the expression of C lineage specific genes (Baugh et al., 2005; Hunter and Kenyon, 1996). Altogether, these observations suggest that a high SKN-1/PAL-1 ratio triggers E specification, while a low SKN-1/PAL-1 ratio induces C specification (Hunter and Kenyon, 1996; Maduro et al., 2005b). Thus, the ectopic expression of *end-1* in *par-4(it47)* embryos could either result from the abnormal stabilization or activity of SKN-1 or from the reduced expression of PAL-1 in the C lineage. Although we could not directly assess SKN-1 activity, our unpublished observations did not reveal any SKN-1 stabilization in *par-4(it47)* embryos. By contrast PAR-4 has previously been shown to be required for PAL-1 accumulation in EMS and P2 cells (Bowerman et al., 1997), suggesting that reduced PAL-1 levels could be the cause of *end-1* expression in the C lineage of *par-4(it47)* embryos.

Importantly, most C cells ectopically expressing *end-1* also in turn activate *elt-2* expression. This observation is consistent with the terminal phenotype of *par-4* embryos, which exhibit an excess of differentiated intestinal cells. The number of cells that eventually express *elt-2* is however variable. Previous studies have shown that robust *elt*-2 expression relies on robust END-1 and END-3 levels and that variability of their expression leads to stochastic *elt*-2 expression (Choi et al., 2017; Raj et al., 2010). It is thus likely that variations in either the levels or the timing of *end-1* expression in the C lineage of *par-4* embryos result in variable *elt-2* expression. Notably, we also observe that some E descendants did not express *elt-2* in *par-4* embryos. These results are reminiscent of previous studies showing that PAR-4 is required for the formation of differentiated intestinal cells (Kemphues et al., 1988; Morton et al., 1992). They suggest that PAR-4 does not only prevent *end-1* expression in the C lineage but may also ensure its robust expression in the E lineage.

At the time when C descendants start to express *end-1* and then *elt-2*, they are located in the vicinity of E descendants but need to rearrange and incorporate to E descendants to form a continuous intestine. As previously described in *cdc-25.1(gof)* mutants (Choi et al., 2017), supernumerary cells eventually accommodate to form rings with more than two cells surrounding the lumen. We cannot exclude that they also occasionally form additional intestinal rings. The presence of lumen deformations is linked to the presence of supernumerary enterocytes. It is nevertheless intriguing that they become apparent only at the 3-4-fold stages. One possible explanation is that this is due to the progressive and late incorporation of supernumerary cells to form intestinal rings containing more than two cells. Alternatively, supernumerary cells may already be incorporated into intestinal rings at the 2-fold stage and induce weak lumen deformations. Those small and early lumen deformations may then be amplified and become detectable as the lumen grows. The presence of additional CeAJs in both 2 and 3-4-fold embryos support this latter hypothesis.

In conclusion, our work reveals a novel function of PAR-4 in regulating intestinal specification. While previous work showed that PAR-4 is required to induce intestinal differentiation in the E lineage, our results demonstrate that it also prevents the C blastomere from producing intestinal cells. Similarly to supernumerary cells arising from additional divisions inside the E lineage, extra intestinal cells resulting from abnormal specification have the astonishing ability to incorporate within the intestinal epithelium, eventually forming a continuous, albeit deformed, intestinal lumen.

## Materials and Methods

### Worm strains

Strains were grown on agar plates containing NGM growth media and seeded with *E. coli* (OP50 strain). Worms were maintained at 20°C, except strains carrying the *par-4(it47)* thermosensitive mutation, which were kept at 15°C and shifted at restrictive temperature (25°C) during the course of our experiments. The strains used in this study are listed in Table S1 and the temperature conditions used for each experiment are detailed in Table S2.

### New CRISPR strains

CRISPR-CAS9-genome edited mNG::PAR-4, PAR-4::mKate2::AID and LET-413c::mNG were generated at the “Biologie de Caenorhabditis elegans” facility (Université Lyon 1, UMS3421, Lyon, France). mNG::PAR-4 was obtained by tagging the two longest isoforms of PAR-4 (PAR-4a and PAR-4c) with mNeonGreen at their common N-terminus. PAR-4::mKate2::AID was obtained by inserting a GASGASGAS linker, mKate2 and an Auxin Inducible Degron (AID, MPKDPAKPPAKAQVVGWPPVRSYRKNVMVSCQKSSGGPEAAAFVK) (Zhang et al., 2015) at the N-terminus of the short PAR-4 isoform (PAR-4b). This insertion also tags the two PAR-4 long isoforms (insertion after Met142 in PAR-4a and PAR-4c). LET-413c::mNG was obtained by fusing mNeonGreen to the C-terminus of the LET-413c isoform; a GASGASGAS linker was inserted between LET-413 and mNG.

### Auxin induced degradation

Intestine specific degradation of PAR-4 was induced by incubating at 20°C embryos expressing PAR-4::mKate2::AID and the auxin receptor TIR-1 under the control of the *elt-2* promoter in 5 mM IAA-AM (acetoxymethyl indole-3-acetic acid, a cell permeable form of auxin which is able to trigger degradation in embryos, (Negishi et al., 2019)). Control experiments showed that PAR-4 was absent from intestinal cells after 1h15 auxin treatment. The number of intestinal cells in embryos depleted of PAR-4 was assessed after 6h auxin treatment.

### Microscopy and time-lapse recordings

For standard and superresolution confocal microscopy, embryos were mounted on 2% agarose pads in a drop of M9 medium. Standard confocal images were acquired with a SP8 confocal microscope (Leica) equipped with a HC Plan-Apo 63×, 1.4 NA objective and the LAS AF software. Z-stacks were acquired with 400 nm steps. Superresolution images were acquired with a LSM 880 microscope (Zeiss), equipped with a Plan-Apo 63×, 1.4 NA objective and the Zen Black software. The Airy Scan module was used to obtain superresolution images. Z-stacks were acquired with 180 nm steps (optimal step size).

For lineage experiments, adult hermaphrodites were dissected in M9 medium. Embryos were then transferred to a 4 μL drop of M9 medium on a 35 mm glass bottom dish (poly-D-lysine coated, 14 mm microwell, No 1.5 coverglass, MatTek). Excess of M9 was then removed to ensure adhesion of embryos to poly-D-Lysine and 2 mL of mineral oil (Light Oil (neat), Sigma) was used to cover the drop of M9. Recordings were performed on a Leica DMi8 spinning disc microscope equipped with a HCX Plan-Apo 63×, 1.4 NA objective and a Roper Evolve EMCCD camera. The setup was controlled by the Inscoper Imaging Suite. Embryos were maintained at 25°C during recordings. Images were acquired at 3 or 5 minutes intervals (Fig.5B and Fig.6D) or every 5 minutes during the first two hours of recordings and then every 10 minutes (Fig.6A,E-H, Fig.S5). Z-stacks were acquired with 400 nm steps.

Temperature conditions used for each experiment are detailed in Table S2.

### Transmission electron microscopy

Control and mutant *C. elegans* embryos (at 3-4 fold stage) were shifted at 25°C two hours before fixation. They were fixed by high pressure freezing with the EMPACT-2 system (Leica Microsystems). Cryosubstitution (FS) was done in anhydrous acetone containing 1% OsO4, 0.5% glutaraldehyde and 0.25% uranyl acetate for 60 h in an FS system (AFS-2; Leica Microsystems). The embryos were embedded in an Epon-Araldite mix (EMS hard formula). Adhesive frames were used (11560294 GENE-FRAME 65L, Thermo Fisher Scientific) for flat embedding, as previously described (Bidaud-Meynard et al., 2019). Ultrathin sections (70 nm thickness) were cut on an ultramicrotome (UC7; Leica Microsystems) and collected on formvar coated slot grids (FCF2010-CU; EMS). Each sample was sectioned at 7-10 different places with ≥5 μm between each grid, to ensure that different gut cells were observed. TEM grids were then observed using a JEM-1400 TEM (JEOL) operated at 120 kV, equipped with a Gatan Orius SC1000 camera and piloted by the Digital Micrograph 3.5 program (Gatan).

### Image analysis and quantifications

Images were assembled for illustration using the Fiji and Inkscape 1.1 softwares. All embryos are oriented with the anterior pole to the left. Images obtained with the Airy Scan module were processed to enhance resolution with the ZenBlack software. Max intensity z-projections, orthogonal views and 3D reconstructions were obtained with Fiji. In lineage experiments, there are very strong differences in signal intensities between the *end-1p*::GFP::H2B and *elt-2p*::NLS::GFP::LacZ reporters, with the *elt-2* signal being much stronger than the *end-1* signal. The signal intensities on images from lineage movies were thus adjusted to enable proper visualization of intestinal nuclei at all the embryonic stages observed.

TEM micrographs were analyzed using Fiji. Microvilli length was measured from the tip to the point where their base intersected with the apical pole. Microvilli width was measured at mid-height. Microvilli density was defined as the number of microvilli over 1 μm of lumen perimeter. Junction length correspond to the length of the electron-dense structure.

### Statistical analysis

Statistical analysis and graphic representations were performed with the Graphpad Prism 8.0.1 software. Details of the statistical tests used are indicated in each figure legend.

## Acknowledgments

We thank M. Boxem, B. Grant, M. Soto and the *Caenorhabitis* Genetics Center (funded by the National Institute of Health Office of Research Infrastructure Programs, P40 OD010440) for providing worm strains. We also thank the Biology of *Caenorhabditis elegans* Facility (Université Lyon 1, UMS3421) for generating CRISPR strains. We thank Anaëlle Raoul who helped with the characterization of intestinal cell number in *par-4* mutants as well as the Michaux and Pecreaux labs for helpful discussions. Imaging was performed at the Microscopy Rennes imaging Center (MRiC Photonics and TEM, Biosit, Rennes, France), a member of the national infrastructure France-BioImaging supported by the French National Research Agency (ANR-10-INBS-04).

## Author contributions

Conceptualization: F.D., G.M., A.P.; Methodology: F.D., O.N., A.P.; Validation: F.D., O.N., G.M., A.P.; Formal analysis: F.D., O.N., A.P.; Investigation: F.D., O.N., A.P.; Writing – original draft preparation: F.D., A.P.; Writing – Review and editing: F.D., O.N., G.M., A.P.; Vizualization: F.D., O.N.; Supervision: G.M., A.P.; Project administration: G.M.; Funding Acquisition: F.D., G.M., A.P.

## Figure legends

**Figure S1:**
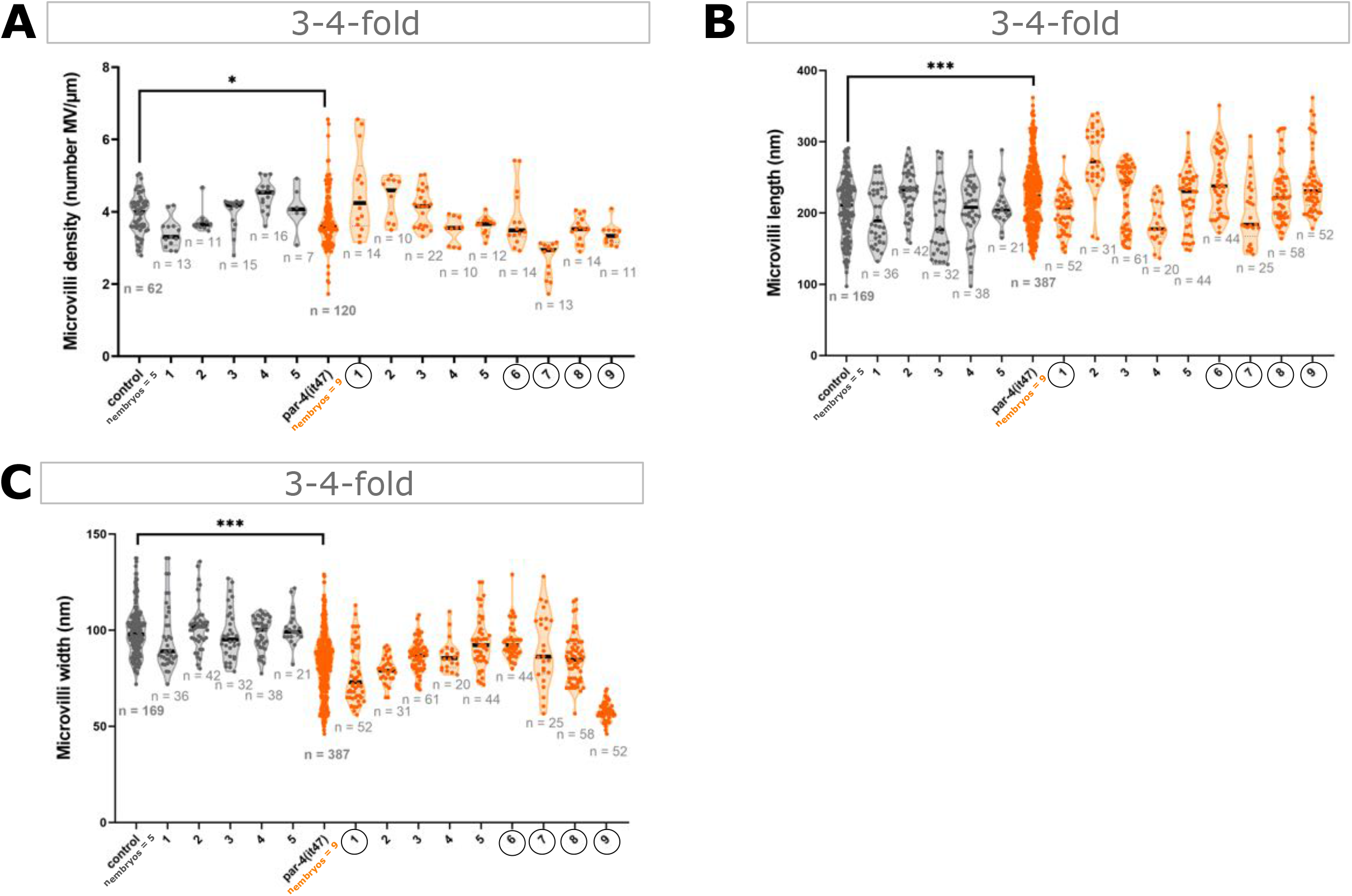
PAR-4 loss-of-function induces mild microvilli defects. **(A-C)** Quantification of microvilli density (A), length (B) and width (C) in 3-4-fold control and *par-4(it47)* embryos from TEM images. Microvilli density corresponds to the number of microvilli per μm. Values were obtained from 5 control and 9 *par-4(it47)* embryos, measuring density in more than 7 areas and microvilli characteristics for more than 20 microvilli per embryo. Violin plots show all individual values (dots) and the mean (bold black line). Control and *par-4(it47)* plots on the left were obtained by pulling the data of all embryos, other plots correspond to the measurements made for each single embryo. n indicates the number of either areas or microvilli measured. Samples for which sections with more than 5 CeAJs and more than 4 intestinal cells per intestinal ring were observed are circled. *P<0.05, ***P<0.001. Statistical significance was calculated using a Mann-Whitney test.

**Figure S2:**
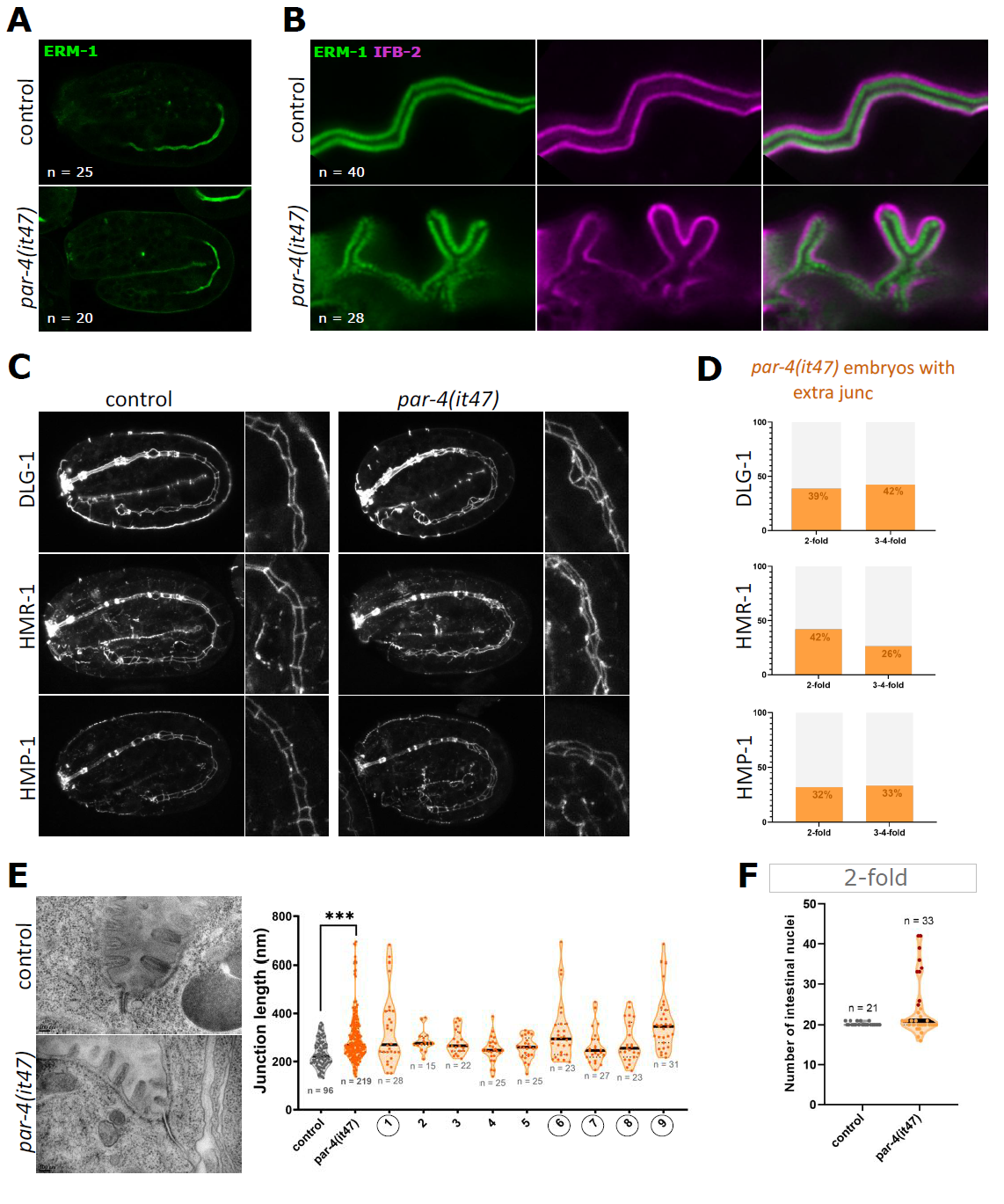
PAR-4 controls lumen morphology, apical junction number and junction length. **(A)** Maximum Z-projections from confocal images of 2-fold control and *par-4(it47)* embryos expressing ERM-1::mNG. Scale bar: 5 μm. **(B)** SR confocal images of 3-4-fold control and *par-4(it47)* embryos co-expressing ERM-1::mNG (green) and IFB-2::wScarlet (magenta). Arrowheads indicate apical membrane deformations. Scale bar: 1 μm. **(C)** Maximum Z-projections from confocal images of 2-fold control and *par-4(it47)* embryos expressing DLG-1::mNG, HMR-1::GFP or HMP-1::GFP. Zoomed-in images show high magnification of apical junctions. Scale bar: 5 μm. **(D)** Quantification of the percentage of 2-fold and 3-4-fold *par-4(it47)* embryos with additional CeAJs in the three different strains. **(E)** TEM images of transversal sections of 3-4-fold control and *par-4(it47)* embryos and quantification of junction length. CeAJs are colored in magenta. Scale bar: 100 nm. Values were obtained from 5 control and 9 *par-4(it47)* embryos, measuring more than 15 junctions for each embryo. Violin plots show all individual values (dots) and the mean (bold black line). Control and *par-4(it47)* plots on the left were obtained by pulling the data of all embryos, other plots correspond to the measurements made for each single *par-4(it47)* embryo. n indicates the number of junctions measured. Samples for which sections with more than 5 CeAJs and more than 4 intestinal cells per intestinal ring were observed are circled. Violin plots show all individual values (dots) and the mean (bold black line). ***P<0.001. Statistical significance was calculated using a Mann-Whitney test. **(F)** Quantification of intestinal nuclei number in 2-fold control and *par-4(it47)* embryos at permissive temperature (15°C). Violin plots show all individual values (dots) and the mean (bold black line).

**Figure S3:**
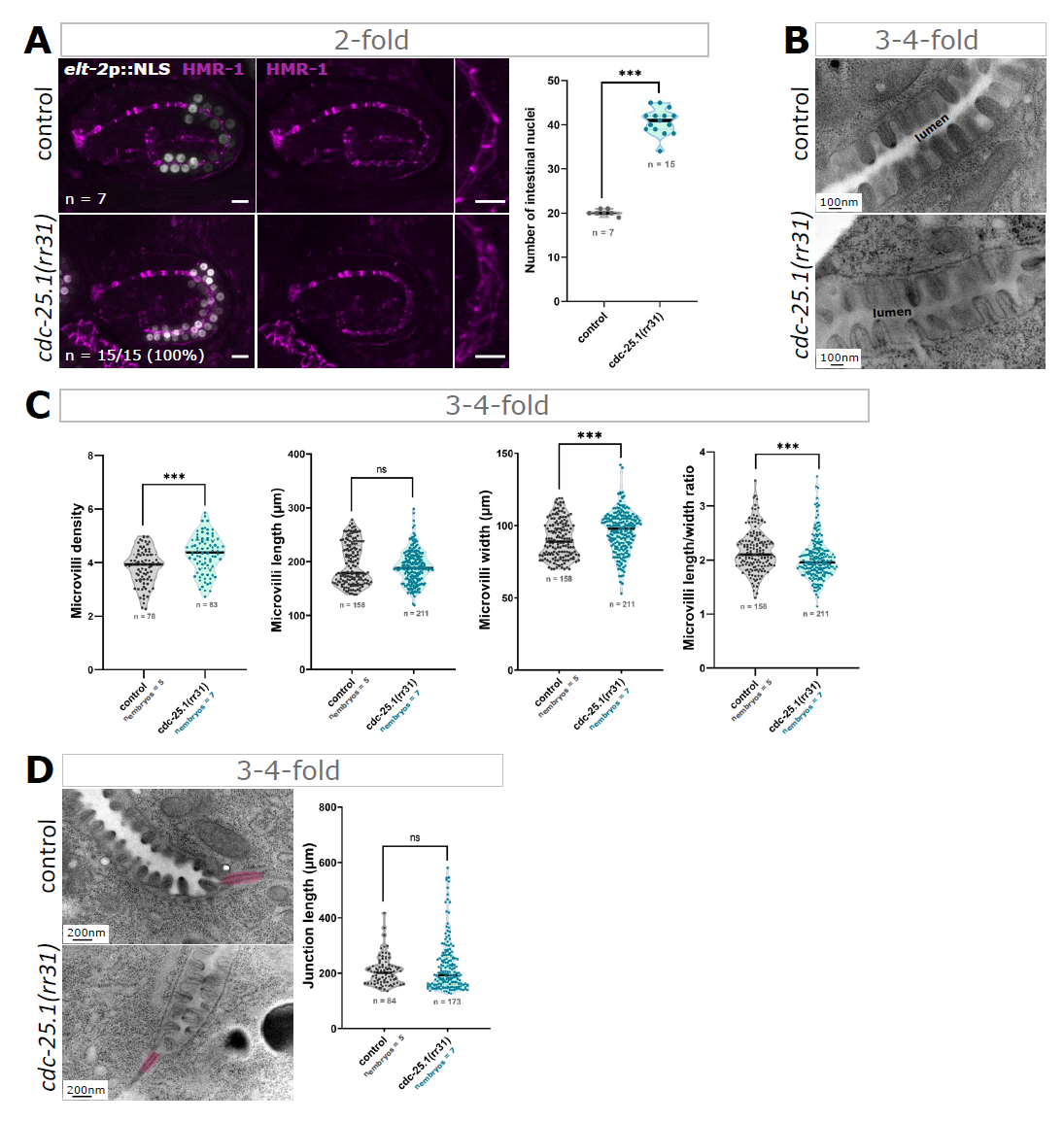
CDC-25.1 gain-of-function phenotype is not completely similar to the phenotype induced by PAR-4 loss-of-function. **(A)** Maximum Z-projections from confocal images of 2-fold control and *cdc-25.1(rr31)* embryos co-expressing *elt-2p*::NLS::GFP::LacZ and HMR-1::mKate2 and quantification of the number of intestinal nuclei in those embryos. Zoomed-in images showing HMR-1::mKate2 marker. Scale bar: 5 μm. Violin plots show all individual values, plus the mean (bold black line). ***P<0.001. Statistical significance was calculated using a Student’s *t*-test. **(B)** TEM images of transversal sections of the brush border in 3-4-fold control and *cdc-25.1(rr31)* embryos. Scale bar: 100 nm. **(C)** Quantification of microvilli density, length, width and length/width ratio in 3-4-fold control and *cdc-25.1(rr31)* embryos from TEM images. Microvilli density corresponds to the number of microvilli per μm. **(D)** TEM images of transversal sections of 3-4-fold control and *cdc-25.1(rr31)* embryos and quantification of junction length. CeAJs are colored in magenta. Scale bar: 200 nm. In (C-D) values were obtained from 5 control and 7 *cdc-25.1(rr31)* embryos, measuring density in more than 10 areas, microvilli characteristics for more than 19 microvilli and junction length for more than 11 junctions per embryo. Violin plots show all individual values (dots) and the mean (bold black line). They were obtained by pulling the data of all embryos. n indicates the total number of either areas, microvilli or junctions measured. ns: non-significant; P>0.05, ***P<0.001. Statistical significance was calculated using a Student’s *t*-test (for microvilli density) or a Mann-Whitney test (for other parameters).

**Figure S4:**
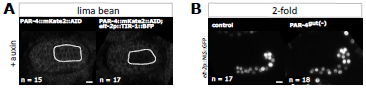
PAR-4 degradation in enterocytes does not affect intestinal cell number. **(A)** Confocal images of E16 embryos expressing only PAR-4::mKate2::AID or both PAR-4::mKate2::AID and *elt-2p*::TIR-1::BFP after 1h15 auxin treatment. In embryos co-expressing these transgenes, PAR-4 is specifically degraded in intestinal cells (surrounded with dashed line). These embryos are called PAR-4^gut(-)^ embryos. Scale bar: 5μm. **(B)** Maximum Z-projections from confocal images of 2-fold embryos expressing *elt-2p*::NLS::GFP::LacZ in a control or a PAR-4^gut(-)^ background after 6h auxin treatment. Scale bar: 5μm.

**Figure S5:**
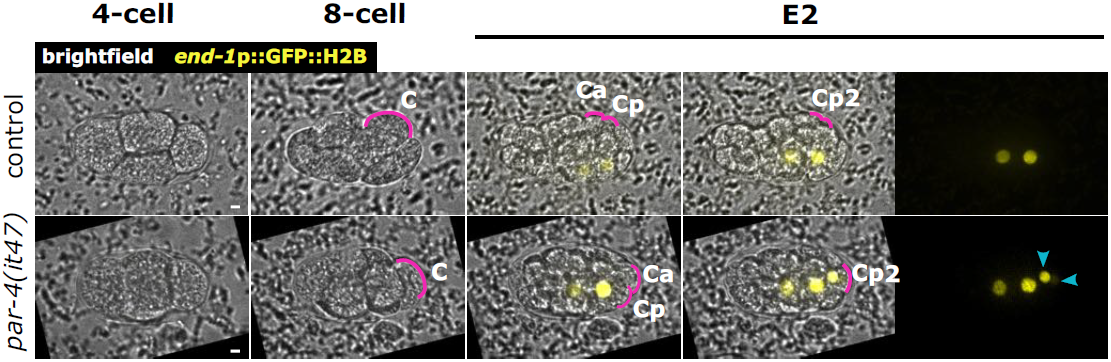
PAR-4 prevents expression of END-1 in the C lineage. Cell lineage movies in control and *par-4(it47)* embryos expressing *end-1*p::GFP::H2B and *elt-2p*::NLS::GFP::LacZ. Embryo morphology is shown through the brightfield channel. The position of C, Ca, Cp and Cpp/Cpp (noted Cp2) blastomeres is indicated with the magenta line. Images of *end-1* expressing nuclei (yellow) are superposed with brightfield images (left panels) or shown as maximum Z-projections (right panels). Blue arrowheads indicate nuclei from Cpa and Cpp nuclei expressing *end-1. end-1* is also expressed in two other nuclei, which correspond to Ea and Ep. Scale bar: 5μm.

**Movie 1: Dynamic behaviour of mNG::PAR-4 foci in polarizing enterocytes**. Large PAR-4 foci move in the vicinity of the apical membrane while small foci move along the lateral membranes. Images were acquired every 15 sec and movie is played at 10 frames per second.

**Table S1:**
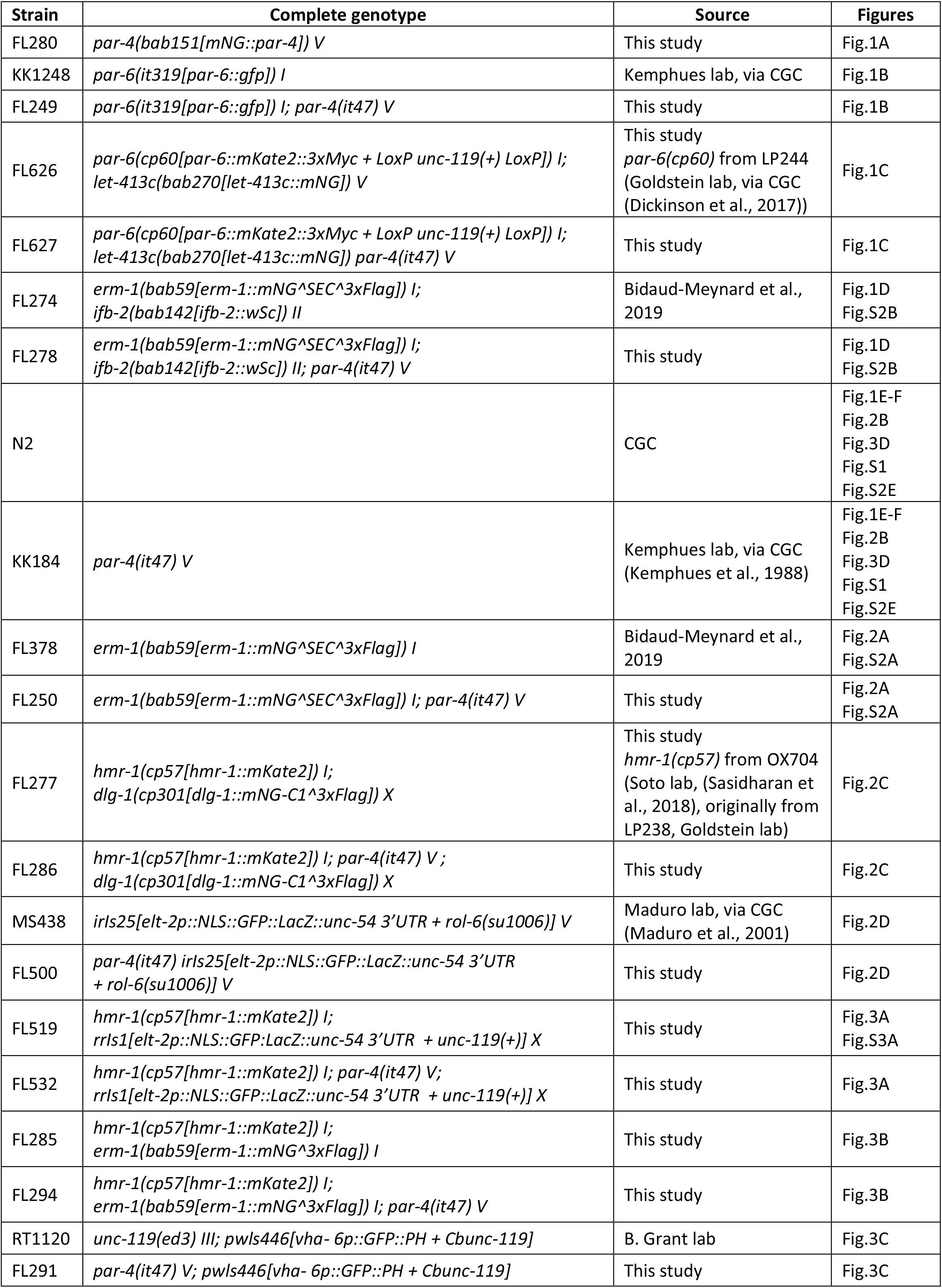
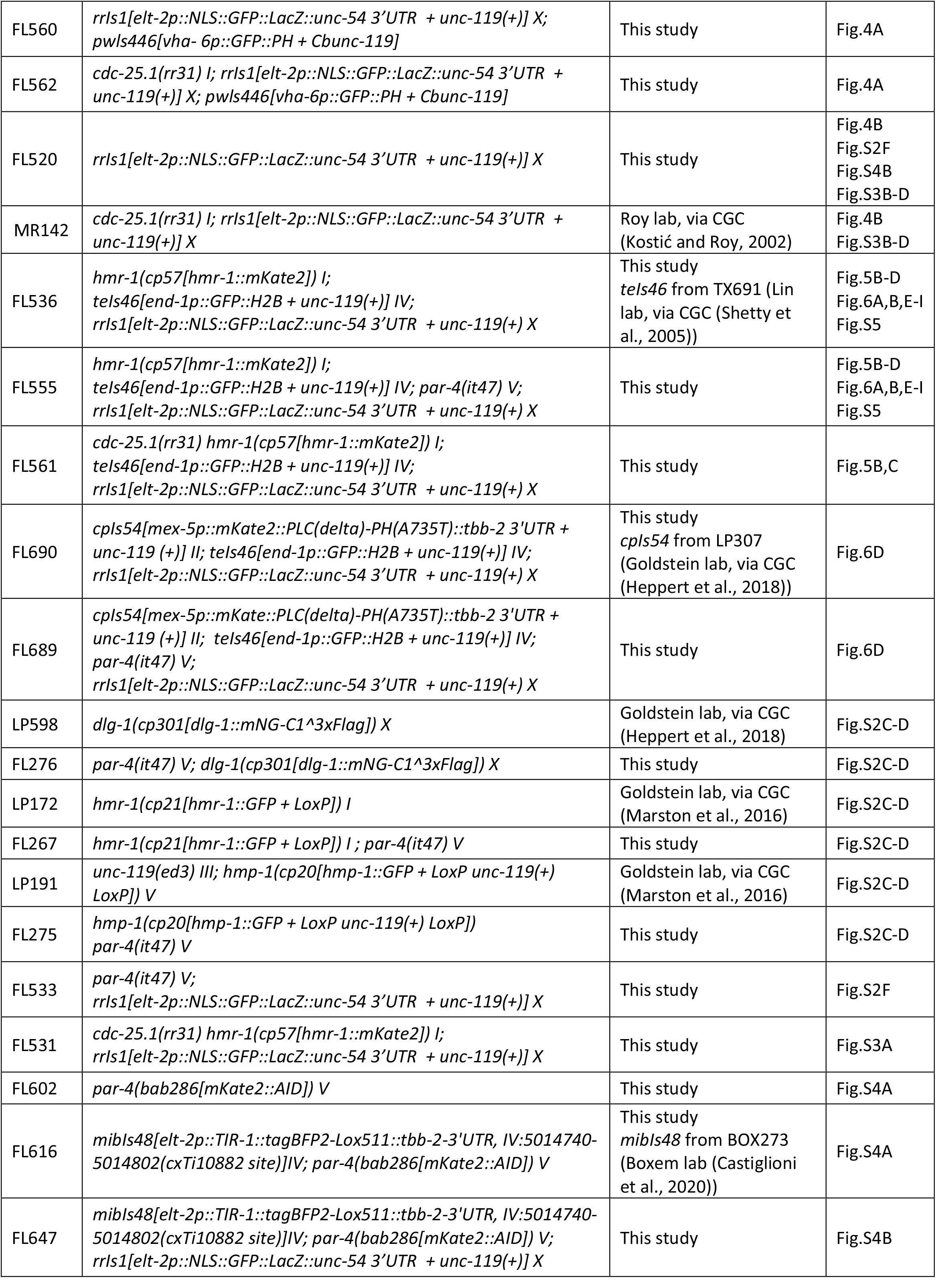
*C. elegans* strains used in this study.

**Table S2:** Summary of experimental conditions used.

| **Temperature conditions used before and during experiments** | **Figures** |
| --- | --- |
| Strains grown at 15°C, no temperature shift | Fig.S2F |
| Strains grown at 20°C, no temperature shift<br>For auxin experiments, incubation at 20°C | Fig.1A, Fig.4A<br>Fig.S3A, Fig.S4 |
| Strains shifted at 25°C 2h before starting imaging or fixation (TEM),<br>maintained at 25°C during time-lapse recordings | Fig.1B-F, Fig.2, Fig.3A-B,D, Fig.4B, Fig.5D<br>Fig.S1, Fig.S2A-E, Fig.S3B-D |
| Strains shifted at 25°C just at the beginning of imaging,<br>maintained at 25°C during time-lapse recordings. | Fig.5B-C |
| Strains shifted at 25°C 1h before starting imaging,<br>maintained at 25°C during time-lapse recordings | Fig.6,<br>Fig.S5 |
| Strains shifted at 25°C 3h before starting imaging,<br>maintained at 25°C during time-lapse recordings | Fig.3C |

